# Large language models possess some ecological knowledge, but how much?

**DOI:** 10.1101/2025.02.10.637097

**Authors:** Filip Dorm, Joseph Millard, Drew Purves, Michael Harfoot, Oisin Mac Aodha

## Abstract

Large Language Models (LLMs) have shown remarkable capabilities in question answering across various domains, yet their effectiveness in ecological knowledge remains underexplored. Understanding their potential to recall and synthesize ecological information is crucial as AI tools become increasingly integrated into scientific workflows. Here, we assess the ecological knowledge of two LLMs, `Gemini 1.5 Pro` and `GPT-4o`, across a suite of ecologically focused tasks. These tasks evaluate an LLM’s ability to predict species presence, generate range maps, list critically endangered species, classify threats, and estimate species traits. We introduce a new benchmark dataset to quantify LLM performance against expert-derived data. While the LLMs tested outperform naive baselines, achieving around 20 percentage points higher accuracy in species presence prediction, they reach only a third of the mean F1 score for range map generation and improve threat classification by just around 10 points over random guessing. These results highlight both the promise and challenges of applying LLMs in ecology. Our findings suggest that domain-specific fine-tuning is necessary to improve ecological knowledge in LLMs. By providing a repeatable evaluation framework, our benchmark dataset will facilitate future research in this area, helping to refine AI applications for ecological science.

## 1. Introduction

The biodiversity crisis is one of the defining challenges of our time, making data collation on biodiversity change increasingly important. As species are being lost at an unprecedented rate [1], ecologists are urgently working to address persistent gaps in our understanding of global biodiversity patterns [2]. Traditionally, biodiversity research has relied on manual curation and fragmented datasets, limiting both coverage and scalability. These persistent knowledge gaps and methodological limitations have prompted growing interest in new approaches to biodiversity data collection and analysis. In particular, recent efforts have focused on automating information extraction from the scientific literature, using tools like text mining to overcome the challenges of manual synthesis [3, 4].

Large Language Models (LLMs) have emerged as a powerful technology, demonstrating strong performance across a wide range of tasks, including coding, mathematics, and language understanding [5, 6, 7, 8]. These models, trained on massive corpora using significant computational resources, follow scaling laws that place the development of high-performing LLMs far beyond the reach of most ecological researchers. As a result, researchers in ecology are likely to turn to general-purpose, off-the-shelf models to support a wide range of tasks involving information extraction and synthesis. Despite growing enthusiasm, the baseline capabilities of LLMs in ecology use cases remain largely untested. While benchmarking studies have evaluated model performance in several fields [9], there has yet to be a systematic assessment of how these models perform on ecological knowledge tasks.

One related domain where interest in LLM applications has grown rapidly is geospatial analysis [10, 11, 12, 13, 14, 15]. Particularly relevant to ecological applications is GPT4GEO [16], which includes evaluation tasks such as estimating the spatial distribution of species and bird migratory routes. While GPT4GEO represents an important step toward connecting LLMs and ecology, it focuses mainly on spatial inference and qualitative assessment. These geospatial LLM analyses find that models tend to perform better when locations are described using place names, likely due to the alignment with the textual format of their training data [14]. More broadly, previous work has bridged geospatial analysis and ecology, with a focus on text retrieval for modeling the distribution of species [17, 18, 19]. These developments underscore growing interest in ecological applications of LLMs, not only for spatial inference, but increasingly for tasks like retrieving and synthesizing ecological evidence. Together, these studies reveal progress toward applying LLMs in ecology, but they leave open how comprehensively models capture ecological knowledge, a gap we aim to address.

Ecological applications of LLMs are new, but developing quickly, particularly in retrieving and synthesizing text and data. Previous research demonstrates LLMs’ ability to assist with tasks in ecology such as extracting information from scientific literature and coding [20, 21, 22, 23, 24, 25, 26]. To address challenges related to erroneous outputs, [27] explored methods for estimating uncertainty in LLM-generated results, helping to mitigate risks in ecological applications. LLMs have also shown promise in debunking sensational wildlife news stories [28] and assessing invasive species distributions based on news reports [29]. Together, these studies highlight the emerging utility of LLMs in ecology, but also show that performance varies with context and task [29, 28].

Here, we present a suite of species-centric evaluation tasks to assess LLMs’ knowledge across multiple areas of ecology. We evaluate two models, `GPT-4o` from OpenAI [30] and `Gemini 1.5 Pro` from Google [31], through a series of five experiments: predicting species presence (Section 2.3.1), generating range maps (Section 2.3.2), listing threatened species (Section 2.3.3), identifying threats (Section 2.3.4), and estimating traits (Section 2.3.5). By establishing this benchmark, we provide a structured means to quantify what ecological knowledge LLMs encode about the natural world, as well as a scalable foundation for future studies targeting more complex ecological reasoning and synthesis.

Previous evaluation of LLMs for predicting trait-related quantities focuses primarily on extracting information from provided datasets and text [32, 33]. Trait estimation has also been explored using Vision-Language Models (VLMs), which enable reasoning over both images and text [34]. In contrast, we evaluate LLMs in a closed-book setting without external databases or retrieval-augmented generation (i.e., where the model autonomously queries external sources such as web APIs at inference), probing what models can do using only their parameters and prompt-provided information (e.g., definitions or other context needed for the task). Together, these tasks establish a reproducible baseline of ecological capability across both recall and application. Studying intrinsic model behavior in this way is important because it highlights what models have learned, informs when external retrieval is actually necessary, and avoids the costs, latency, and potential noise associated with unnecessary external data access [35]. By focusing on closed-book performance, our work extends previous efforts such as GPT4GEO and introduces a scalable benchmark to systematically quantify the ecological capabilities of LLMs.

## 2. Material and Methods

### 2.1. Models

Our experiments consist of querying two commercial LLMs, `Gemini 1.5 Pro` and `GPT-4o`, through their publicly accessible APIs. We focus on `GPT-4o` and `Gemini 1.5 Pro` as they are among the most widely used commercial LLMs at the time of writing and therefore represent a relevant baseline for ecological applications. Our goal is not to be exhaustive, but to establish a repeatable benchmarking framework that can be applied to additional models in future work. While restricting our evaluation to two models means that our conclusions apply only to these rather than to LLMs more broadly, their prominence and accessibility make them appropriate choices for designing and demonstrating the benchmark setup. Both models are used in the period between August and October 2024. We use the following versions of the models: `gemini-1.5-pro-001` and `gpt-4o-2024-05-13`. For both models, the temperature, a hyperparameter controlling the randomness of token selection, is set to zero to minimize variability across runs and improve reproducibility. All other API parameters are left at their default values. Both models default to a maximum of 4096 output tokens. Our tasks are designed such that generated outputs remain within this limit. Importantly, neither `Gemini 1.5 Pro` nor `GPT-4o` retains conversation history when utilizing their APIs [36, 37], which is advantageous for our purposes, as we do not want the model to learn from previous queries.

### 2.2. Datasets

#### Spatial Distribution Data

The species presence and range map tasks are based on expert-derived range maps of 2,418 species from the International Union for Conservation of Nature (IUCN) Red List [38]. This is the same filtered dataset used in existing work for evaluating species distribution models [39, 40]. In this dataset, species ranges are represented as hexagons of size 250 km², based on a H3 hexagon raster [41], each marked to indicate whether the species is present or absent. The IUCN range maps are compiled through expert review and consensus among multiple assessors. While the IUCN does not publish formal inter-assessor agreement statistics, the maps are considered the global reference standard for species’ extents of occurrence. Minor uncertainty or variation among assessors may persist, which we note as a potential but generally small source of benchmark noise.

#### Threat Data

For the threat classification task, we query the IUCN’s API [42] to extract the Red List status of species, countries they reside in, and threats they face. Each record follows the standardized IUCN classification framework, which undergoes review by multiple assessors to ensure consistency, although inter-expert agreement statistics are not publicly reported.

#### Trait Data

For tasks related to predicting numerical traits for species, we utilize two primary data sources. For bird traits, we use AVONET [43], and for mammals, we use COMBINE [44]. Both datasets encompass a wide range of species and contain average measurements of morphological traits. The publicly available AVONET dataset provides species-level average trait measurements rather than raw individual-level data. The authors report that most morphological variance is explained at higher taxonomic levels (order, family, genus), with comparatively low intraspecies variance, supporting the use of species-level averages for comparative analyses. Nevertheless, some traits in certain taxa exhibit higher inter- or intraspecific variance. Accordingly, while our benchmark uses validated expert datasets, both the expert-derived IUCN maps and species-level trait averages represent well-founded but still approximate estimates of biological reality.

#### 2.2.1. Prompting Strategy

For all tasks, we employ a simple prompting strategy: prompts are generally designed to produce outputs in a structured, parseable format (e.g., Python lists and GeoJSONs). In some instances, where it was clear from initial experiments that a model might not naturally produce a structured output, we explicitly guided it within the prompt to adhere to the desired format. Despite these minor interventions, we deliberately avoid extensive prompt engineering or model-specific optimizations, to reduce the risk of overfitting to a particular LLM.

### 2.3. Evaluation Tasks

We design a set of five evaluation tasks to test the ecological information recall of LLMs (Figure 1). These diverse tasks allow us to test an LLM’s knowledge of the geographical distribution of species, their ability to assess threats, and understand species’ size and morphology.

**Figure 1:**
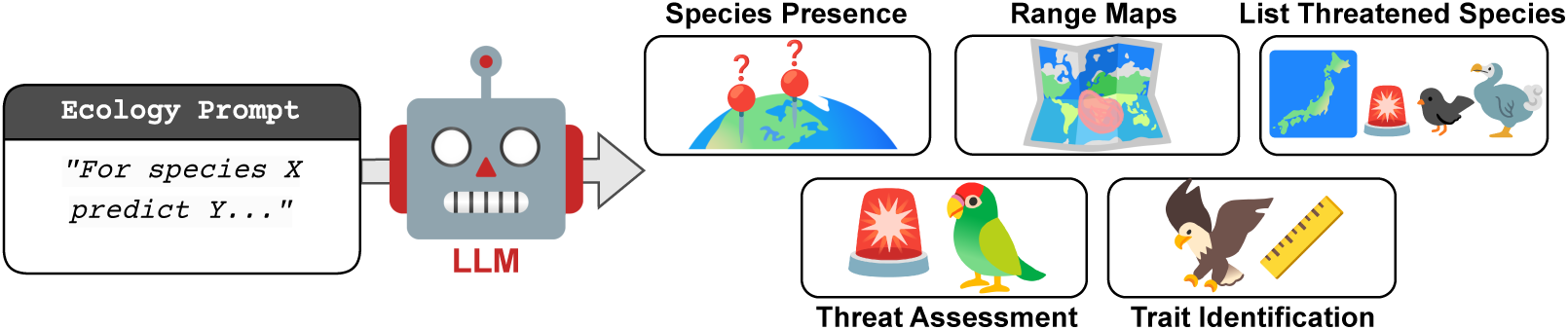
Evaluation Overview. We evaluate existing large language models (LLMs) on a set of five ecological question answering tasks. The tasks require the LLMs to answer questions related to identifying what species are present at a specified location, mapping their ranges, listing threatened species, assessing specific threat levels, and estimating species’ traits.

#### 2.3.1. Task 1: Species Presence

Here we focus on identifying species presence or absence across ten specific locations, aiming to evaluate LLMs’ understanding of species’ geographical distributions. We select 100 species using a stratified random sampling approach to ensure equal representation across four taxonomic groups: 25 amphibians, 25 reptiles, 25 mammals, and 25 birds. This sample size is chosen to balance representation across taxa while remaining computationally feasible, given the cost constraints associated with large-scale LLM inference. Although our design maximizes coverage across major taxonomic groups within available resources, the limited sample size means that small between-model performance differences may remain undetected due to low statistical power. The full list of selected species is available at https://github.com/filipgdorm/eco-llm/blob/main/SPECIES.md.

Each species is paired with ten geographic locations: five where the species is present and five where it is absent. These locations are sampled from IUCN expert-derived range maps. Locations are defined using the centroid coordinates of equal-area hexagonal grid cells, each covering approximately 253 km².

To test the impact of spatial proximity on model performance, two of the five absence locations are intentionally selected to be geographically close to the species’ known range. These difficult absences are drawn from smaller grid cells (253 km²) that fall within larger regional cells of about 12,393 km², which neighbor but do not overlap the known range. This approach ensures that the absence points are near the range boundary while remaining outside it.

We repeat the experiment using country names instead of geographic coordinates (Task 1b). Here, “presences” are defined as countries where the species is known to occur, and “absences” are countries where the species does not occur. The number of presence countries for each species varies, from a minimum of one (for species that inhabit only a single country) to a maximum of five. Absence countries are chosen from neighboring countries where the species is not found. If there are not enough neighboring countries to provide ten absences, random countries are selected as absences. The same prompt structure is used, but instead of specific locations, the countries are listed (see prompt in Figure 2).

**Figure 2:**
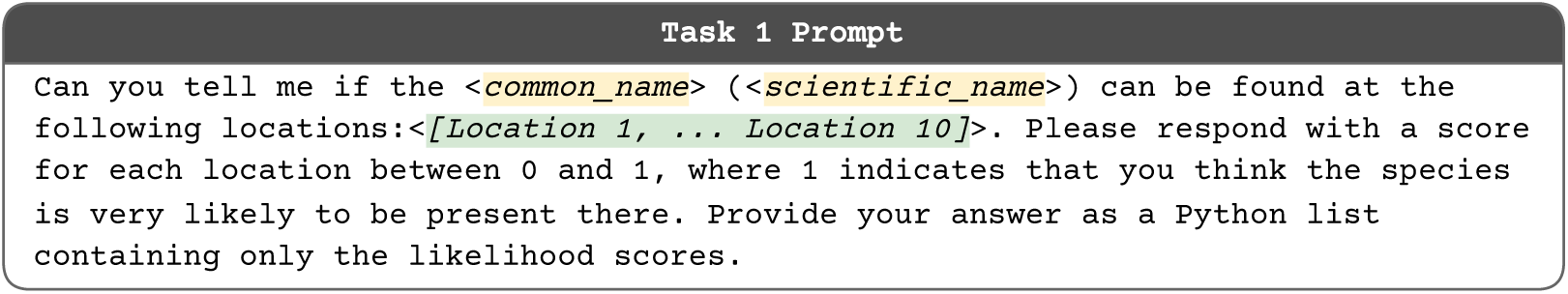
Prompt template for Task 1: Species Presence.

A response is deemed valid if it contains identifiable opening and closing brackets of a Python list, with the content in between being loadable without errors. Evaluation metrics are computed from the valid and parseable responses. As baselines, we include randomly guessing probabilities (i.e., selecting a random number between 0 and 1), always predicting presence (i.e., 1), and always predicting absence (i.e., 0).

To test whether LLM performance on species presence prediction is influenced by model type, taxonomic group, or continent of occurrence, we fit two linear regressions with accuracy as the dependent variable, and LLM type, taxonomic group, and continent presence indicators as predictors. We do this for both Tasks 1a and 1b i.e., coordinates and countries. The continent variables are a set of five binary indicators, each representing whether the species occurs or not in Africa, the Americas, Asia, Europe, or Oceania. Species can occur on multiple continents simultaneously, so these are not mutually exclusive categories. Categorical variables are encoded using treatment coding, with GPT as the reference level for LLM type and Amphibia as the reference level for group name. The models are fit using ordinary least squares regression with the `ols` function using the `statsmodels` Python package [45], and the diagnostic tests are presented in Table C.5 and Table C.6.

#### 2.3.2. Task 2: Range Maps

Here we ask the LLMs to generate range maps for the same 100 species as in Task 1. Assessing LLMs’ geographic awareness through their range map accuracy provides insights into their ability to handle ecological spatial data.

We ask the model to return a GeoJSON string (Figure 3), a plain-text, nested format that is easily parsed and converted into a plottable dataframe [46]. While formats like Shapefiles are common for geospatial data, GeoJSON is more straightforward to handle in this context, since the entire output is contained in a single text string rather than distributed across multiple files, making it easier to parse and analyze directly. The F1 score, measuring how well the GeoJSON-generated range maps match the expert-derived range maps, is computed from this comparison. The F1 score measures how well a model balances precision (the proportion of correctly predicted presences among all predicted presences) and recall (the proportion of correctly predicted presences among all actual presences). Refer to the Appendix B for more details on the F1 score. The GeoJSON is extracted by locating the curly braces {} in the response. The enclosed string is then parsed as a GeoJSON object through Python’s JSON parsing library.

**Figure 3:**
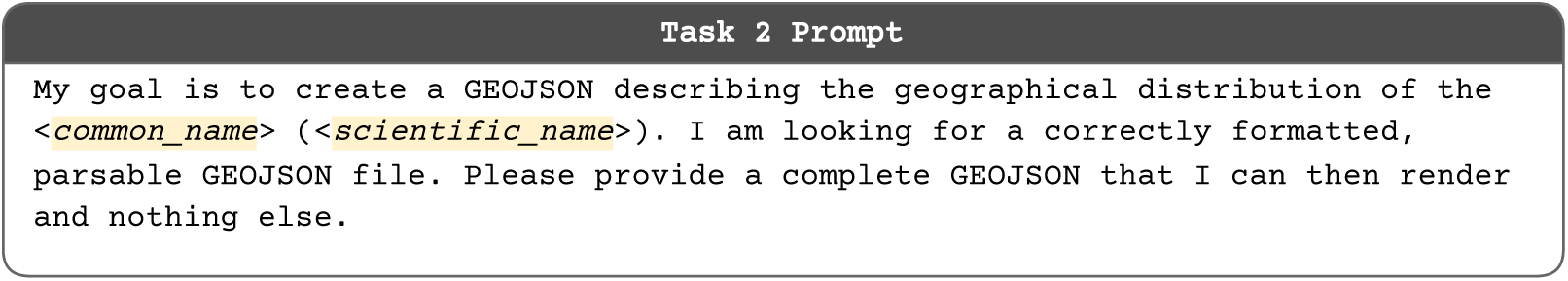
Prompt template for Task 2: Range Maps.

In addition, we also compare our results to a recent deep learning-based species distribution model, SINR [39]. SINR (Spatial Implicit Neural Representations) is a deep learning-based model designed to estimate the distribution of a species from sparse presence-only observations, leveraging large-scale crowdsourced observation data during training. We utilize a coordinate-only version of the SINR model with the *L*_AN-full_ loss function, capped at 1,000 training samples per species and without incorporating any environmental covariates. Predictions are made using either a fixed threshold of 0.1 or a threshold estimated through the LPT-R (Lowest Presence Threshold - Robust) approach as outlined in [40]. The LPT-R method identifies optimal thresholds for binarizing species distribution predictions using only presence data, improving robustness to outliers and offering more accurate range map binarization when only species presence data are available. The mean F1 scores of the range maps produced by SINR and the two binarization methods are computed by comparing to the same IUCN expert range maps as used for the LLMs, ensuring a direct comparison under equivalent data conditions.

The influence of model type, taxonomic group, or continent of occurrence on the F1 score is assessed through linear regression. As in Task 1, the predictors used are LLM type (with levels GPT and Gemini), group name (with levels Amphibia, Aves, Mammalia, and Reptilia), and continent indicators (Africa, the Americas, Asia, Europe, and Oceania). The dependent variable in this case is the F1 score of the range map. The models are fit using ordinary least squares regression with the `ols` function from the `statsmodels` Python package, and the diagnostic tests are presented in Table D.8.

#### 2.3.3. Task 3: List Threatened Species in a Region

Here, we evaluate the ability of LLMs to produce compact summaries of threatened biodiversity by comprehensively listing species that meet specific conservation criteria. Concretely, we focus on species assessed as critically endangered (CR) by the IUCN within a given geographic region (Figure 4). We focus on critically endangered species, since this allows us to quantify the effectiveness of LLMs as tools for identifying species that are the most at risk in a given area. The experiment is intentionally designed to incorporate a geographic element to test how well the two LLMs can synthesize information across domains of ecological knowledge (species, threat status, and location). This approach aligns with real-world conservation use cases, where the geographic scope in assessing conservation status is often constrained to specific regions, such as evaluating the status of species within protected areas, national parks, or even entire countries. The CR category includes species that the IUCN has assessed as having a very high risk of extinction and is limited to those formally assessed, excluding species that may also be at risk but have not yet undergone evaluation.

**Figure 4:**
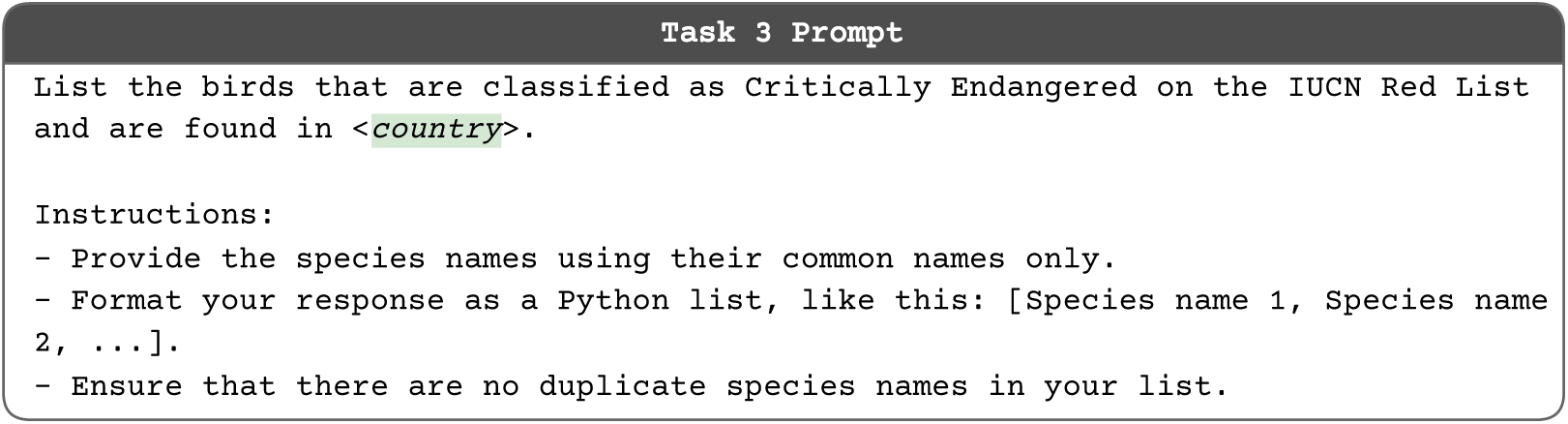
Prompt template for Task 3: Threatened Species.

We choose to limit the task to bird species because including all taxa would be infeasible given response length constraints, and selecting a single well-documented group allows for tractable outputs. Birds are particularly suitable because we have comparatively comprehensive data on them, with large, high-quality datasets available, and their global distributions and conservation assessments are relatively well documented. Also because of the response length constraints, we limit our analysis to common names. In our dataset, common names are shorter on average than scientific binomials, which helps keep model outputs compact. Many practical users also work with common names, so evaluating models in this context is important. Potential ambiguity in naming is handled at evaluation time by leniently mapping model outputs to IUCN records: frequently used alternative common names are treated as correct when they refer to the same taxon, ensuring an unambiguous ground truth.

The task is grounded in political boundaries, meaning that species are assigned to countries based on jurisdictional classifications in the underlying dataset. For instance, both Congo and Lithuania report no critically endangered bird species according to IUCN records, while regions such as Greenland may appear to have no species at all, likely due to their inclusion under Denmark’s jurisdiction. Indonesia has the highest number of CR birds with 30 unique species in that class. Countries without available species data or those not recognized as independent entities by our plotting tool are shown in gray in Figure 9 and are excluded from the summary statistics in Table 3.

#### 2.3.4. Task 4: Threat Assessment

Here, we evaluate the LLMs’ ability to interpret textual threat descriptions and apply ecological knowledge to infer the scope and severity categories defined by the IUCN. In this task, the model is provided with a text description of a threat for a given species (as published by the IUCN), together with the relevant threat code and category definitions (Figure 5). An example of the full information provided to the model is shown in Figure F.15. Using species with known threat classifications, we assess whether the model can reproduce the IUCN scope and severity labels from this text. The species selected for this evaluation collectively span all combinations of severity and scope categories defined by the IUCN, resulting in 100 unique severity–scope pairs. All information is sourced from the IUCN, specifically the Threat Classification documentation [47]. This framing allows us to probe whether LLMs can operationalize provided ecological definitions and make coherent, graded judgments based on text.

**Figure 5:**
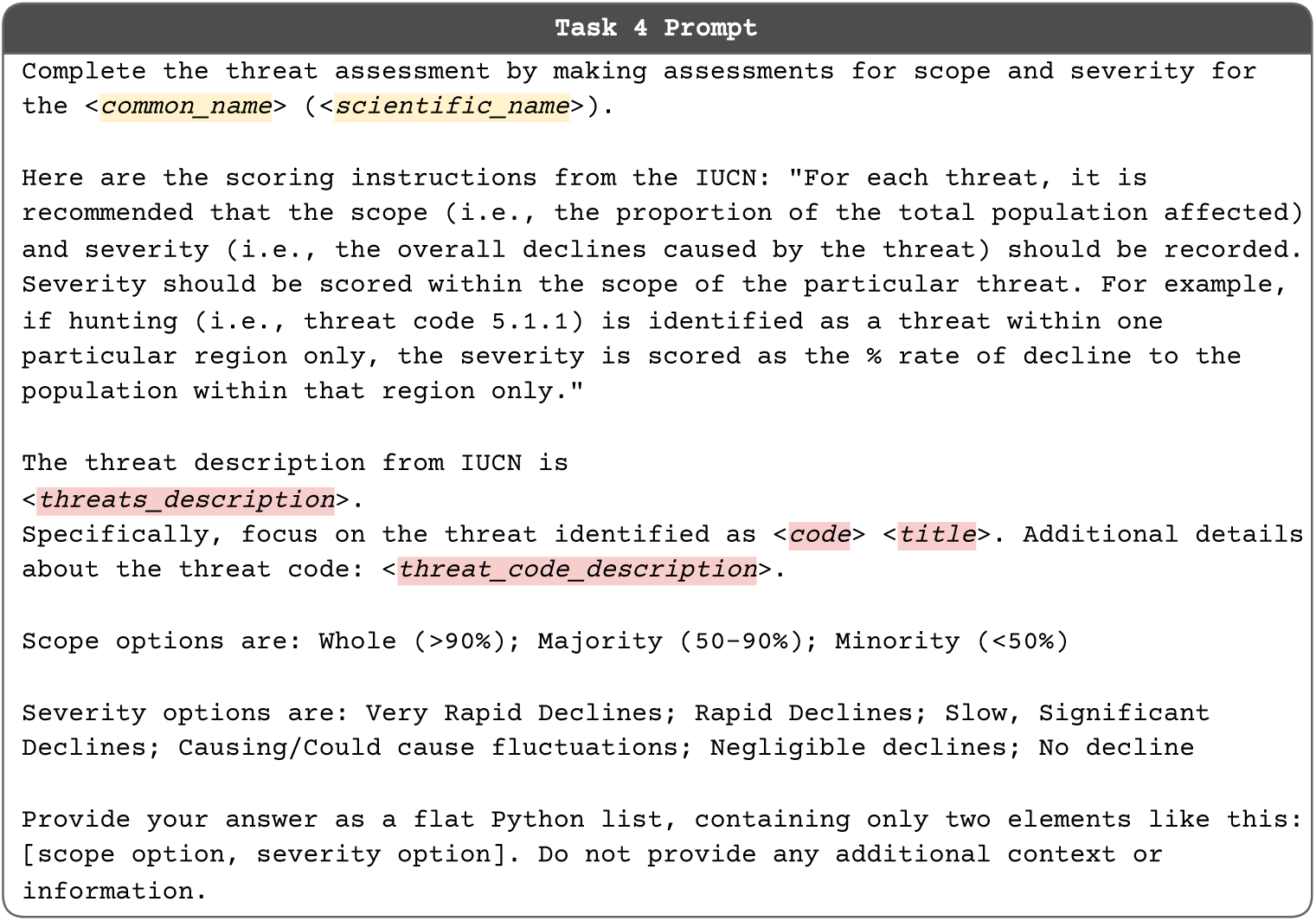
Prompt template for Task 4: Threat Assessment.

To situate this task, it is useful to briefly review how such threat attributes are formalized in conservation practice. Biodiversity is undergoing rapid change due to multiple interacting threats, making accurate characterization of their magnitude and extent essential for conservation. The IUCN Red List provides a standardized framework for this purpose, including categorical classifications of “scope” (the proportion of the population affected) and “severity” (the expected rate of population decline). Scope is represented using the labels “Whole (>90%)”, “Majority (50–90%)”, and “Minority (<50%)” [48], while severity is categorized as “Very Rapid Declines”, “Rapid Declines”, “Slow, Significant Declines”, “Causing/Could Cause Fluctuations”, “Negligible Declines”, or “No Decline” [48]. Although text threat descriptions are often available, these structured labels are frequently omitted, making this task a practical test of whether LLMs can translate unstructured conservation text into standardized assessments.

Because these categories are ordinal rather than nominal, we implement a cost-sensitive performance metric where we account for the severity of misclassifications by awarding partial credit when predicted labels are close to the true labels. For example, predicting “Majority (50–90%)” instead of “Whole (>90%)” in scope classification is considered a smaller error than predicting “Minority (<50%).” We also define a scoring scheme for both scope and severity tasks. For scope, labels (“Minority,” “Majority,” and “Whole”) are mapped to scores (1, 2, and 3). Exact matches score 1.0; adjacent categories score 0.5; distant categories score 0. For severity, six categories are ordered from “No Decline” (1) to “Very Rapid Declines” (6). Matches score 1.0; near matches score 0.75; intermediate 0.5; distant 0.25; and opposites 0.

#### 2.3.5. Task 5: Trait Identification

Here we assess LLMs’ knowledge of traits of different species (Figure 6). We use the AVONET bird [43] and COMBINE mammal [44] trait datasets. From AVONET, we extract the beak, wing, tail length, and mass for 100 randomly sampled bird species. From COMBINE, we select the adult body length, gestation length, max longevity, and adult mass for 100 randomly sampled mammal species. Neither of the datasets makes any distinction between females and males for the traits used, which has an impact on the expected values for some traits. However, given that most species in the sample do not exhibit significant sexual dimorphism, this is not expected to significantly affect the predictive ability of the model.

**Figure 6:**
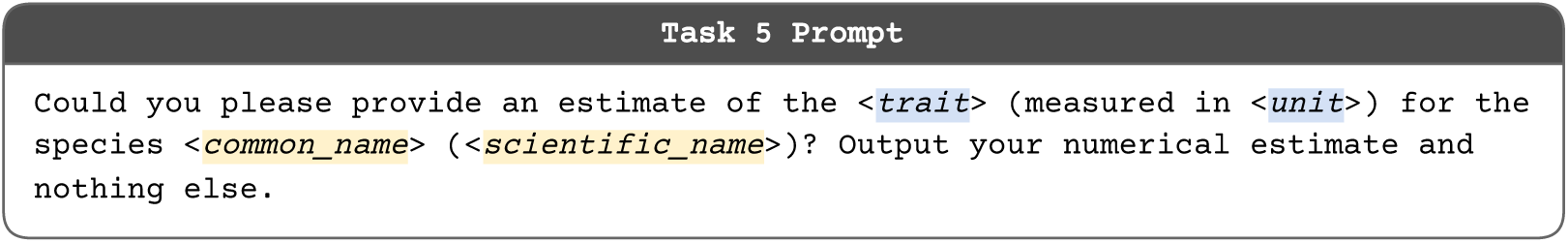
Prompt template for Task 5: Trait Identification.

We ask the models for a numeric answer and specify what unit we want the estimate in, which can both be found in the AVONET and COMBINE datasets (Figure 6). We ask the models to only provide the number, but are lenient to responses that include the unit, e.g., “50 mm”. Wherever units are included in the response, we extract only the numeric substring.

## 3. Results

### 3.1. Task 1: Species Presence

Both `GPT-4o` and `Gemini 1.5 Pro` outperform the random baseline in the presence-absence predictions, particularly when locations are provided as countries rather than precise coordinates (Table 1). While `GPT-4o` shows slightly higher mean accuracy (Table 1, Figure 7), the difference between models is modest. Notably, `Gemini 1.5 Pro` generates more invalid outputs, often due to unparseable formats or because the model refuses to provide an estimate accompanied by a disclaimer. Both models exhibit large standard deviations for their reported accuracies.

**Figure 7:**
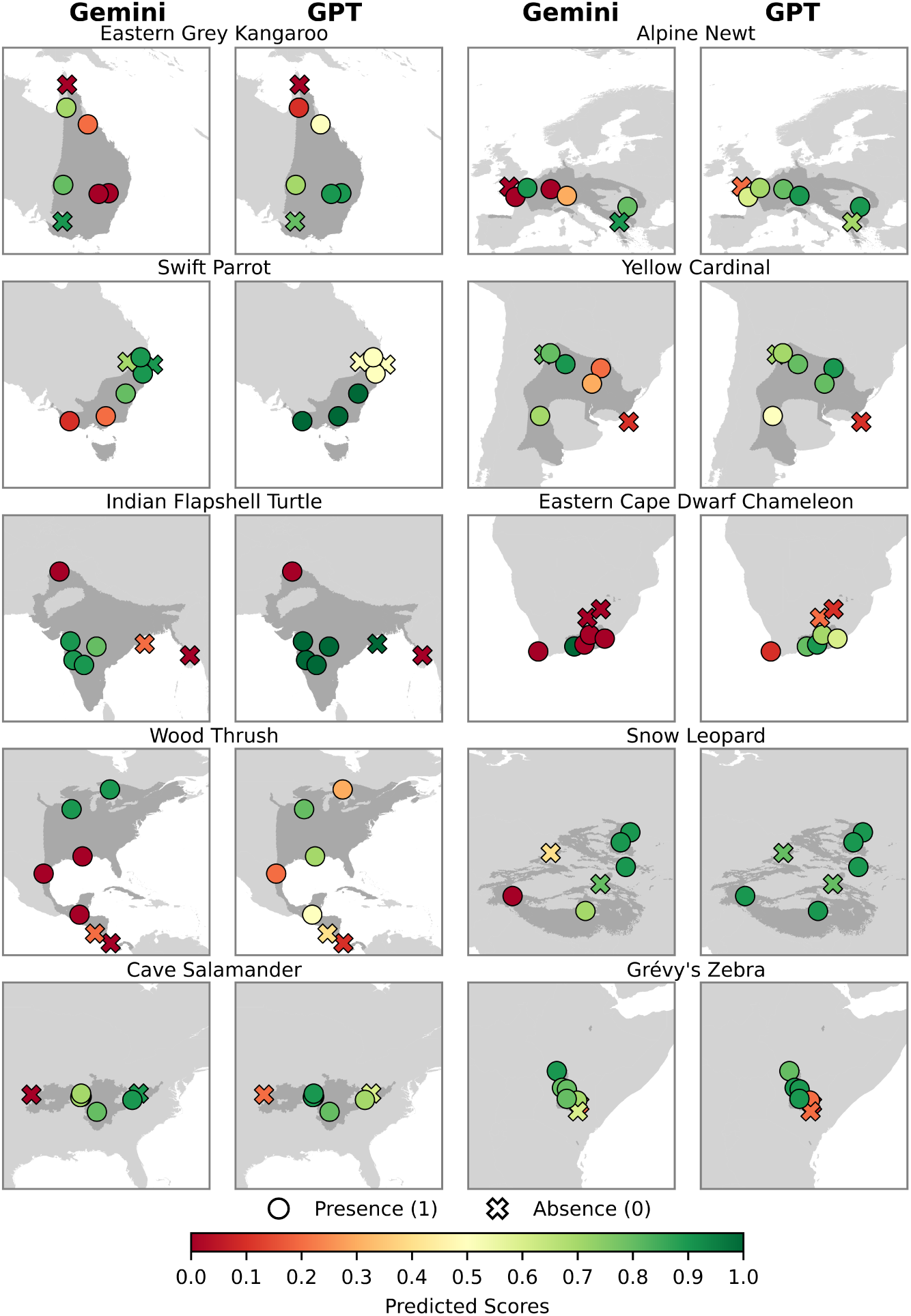
Species Presence. Visualization of selected species spanning all taxonomic groups and IUCN threat categories. Circles show presence locations, crosses show nearby challenging absences. Color indicates predicted presence probability, green is high and close to 1, red is low and close to 0. Dark gray indicates the IUCN range map. A perfect model predicts green for circles and red for crosses.

**Table 1:** Species Presence. . We present mean accuracy (with scores binarized at 0.5) for 100 species, along with the count of unparseable model responses. The “Always Present” baseline (31.7%) is lower than “Always Absent” (68.3%) because most tested species occur in fewer than five countries, so absences dominate. Thus, a naive presence-only model performs worse than an absence-only one.

| Method | Task 1a - Locations |  | Task 1b - Countries |  |
| --- | --- | --- | --- | --- |
|  | Mean Acc. | #Invalid R. | Mean Acc. | #Invalid R. |
| **Gemini 1.5 Pro** | $70.8 \pm 15.7\%$ | 14 | $84.9 \pm 16.1\%$ | 2 |
| **GPT-4o** | **$78.0 \pm 11.9\%$** | 0 | **$87.6 \pm 13.7\%$** | 1 |
| Random Baseline | $51.6 \pm 14.7\%$ | - | $51.3 \pm 12.3\%$ | - |
| Always Absent Baseline | $50.0 \pm 0.0\%$ | - | $68.3 \pm 16.8\%$ | - |
| Always Present Baseline | $50.0 \pm 0.0\%$ | - | $31.7 \pm 16.8\%$ | - |

Further analysis (Appendix C.1) reveals that most prediction errors occur at range boundaries, suggesting that both models can distinguish broad species distributions but struggle with fine-grained spatial limits.

When locations are provided as countries (Task 1b), species from Africa (*p* < 0.001), Asia (*p* = 0.003), and Europe (*p* < 0.001) exhibit substantially lower predicted accuracies than those from Oceania or the Americas (Figure C.11, Table C.6). These geographic patterns are absent in Task 1a, where only Europe (*p* = 0.028) shows a significant difference and accuracy does not deviate meaningfully across other continents (Figure C.11, Table C.5). Accuracy differences across taxonomic groups are minimal in both tasks, with no consistent trend. Model predictions do not vary by LLM type, consistent with the non-significant coefficients in our regression results. Non-significant model differences should be interpreted cautiously, as limited sample size may reduce statistical power.

### 3.2. Task 2: Range Maps

While both `GPT-4o` and `Gemini 1.5 Pro` can generate species range maps that overlap with expert-defined ranges, their outputs do not yet match the accuracy of deep learning-based species distribution models (SDMs) (Table 2). For 77 out of 100 species (`GPT-4o`) and 68 out of 100 species (`Gemini 1.5 Pro`), the predicted ranges intersect the expert-defined ranges, indicating that the models are focusing on the correct geographic regions. `Gemini 1.5 Pro` achieves a higher mean F1 score (24.7% vs. 20.9% for `GPT-4o`) (Table 2), but it also produces more invalid range maps (26 vs. 15). Both models exhibit high variability across species, with standard deviations above 20%.

**Table 2:** Range Maps. . Here we report the mean F1 score of the range maps generated by different LLMs compared with the expert-derived maps. In addition, we also compare to a recent deep learning-based species distribution model SINR [39] (specifically using a coordinate only SINR model with the *L*_AN-full_ loss, capped at 1000 training samples per species and without environmental covariates) using either a fixed threshold of 0.1 or a threshold estimated using the LPT-R approach from [40].

| Method | Mean F1 Score | # Invalid R. |
| --- | --- | --- |
| Gemini 1.5 Pro | $24.7 \pm 22.1\%$ | 26 |
| GPT-4o | $20.9 \pm 21.9\%$ | 15 |
| SINR (threshold 0.1) | $54.9 \pm 22.8\%$ | - |
| SINR (threshold LPT-R) | **$60.7 \pm 17.9\%$** | - |

Qualitative inspection reveals important differences, e.g., `Gemini 1.5 Pro` tends to generate range maps at a finer spatial resolution, while `GPT-4o` often returns coarse, square-like predictions. Among the ten example species shown in Figure 8, only `Gemini 1.5 Pro` achieves F1 scores above 60% for three species (Eastern Grey Kangaroo, Alpine Newt, and Wood Thrush), while `GPT-4o` reaches this level for only one (Eastern Grey Kangaroo). However, even these best cases still exhibit low spatial resolution. A typical failure mode for both models involves generating excessively long lists of co-ordinates that exceed response limits or become incoherent, as seen in the Snow Leopard example (Figure 8), where `GPT-4o` fails to generate a range map.

**Figure 8:**
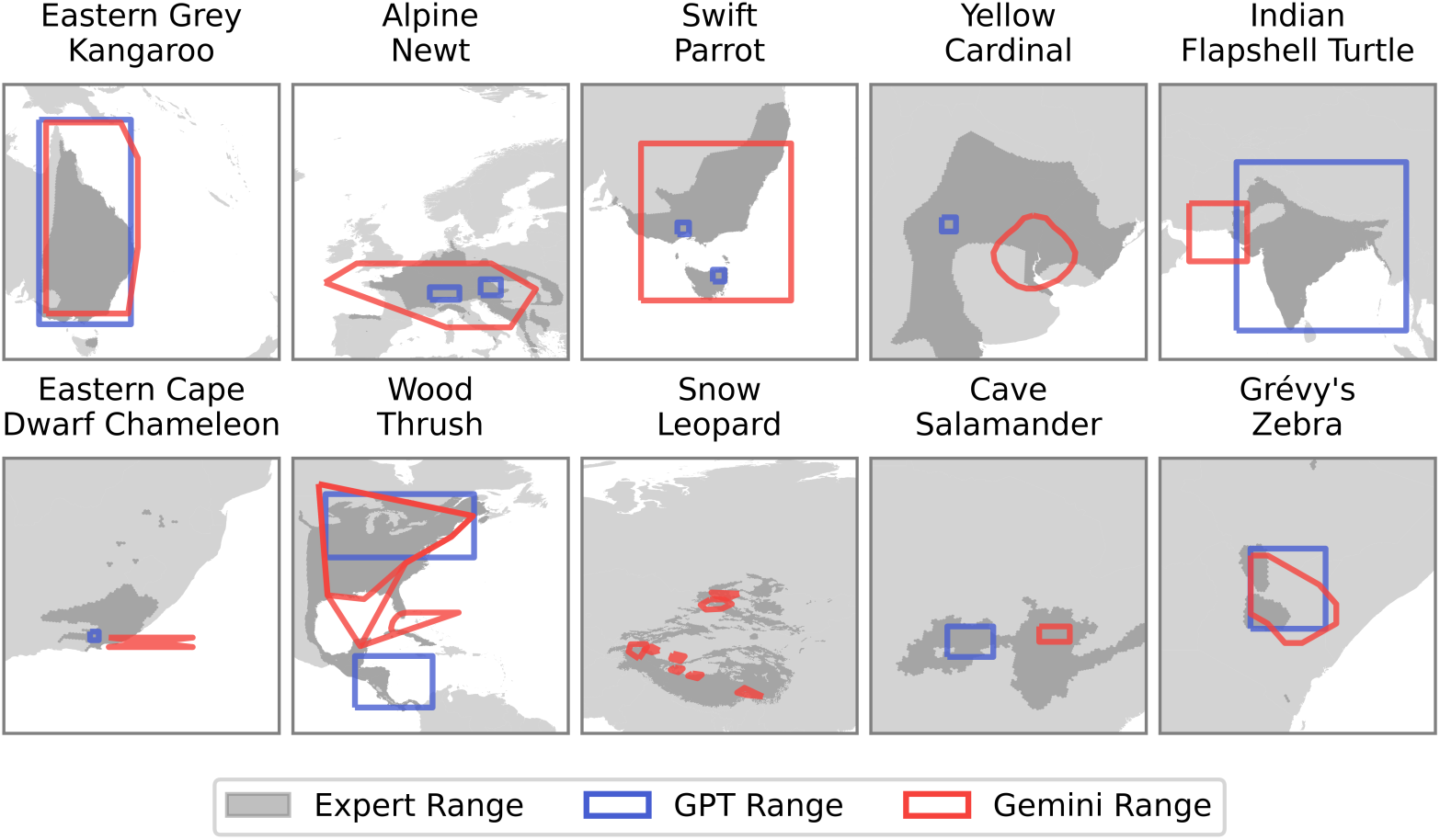
Task 2: Range Maps. Here we display range maps for a subset of species generated by `Gemini 1.5 Pro` (red) and `GPT-4o` (blue) as well as the expert-derived range maps from IUCN (dark gray) we use for evaluation.

We initially explored prompting the models to generate range maps by requesting approximately 50 border coordinates, following the approach in [16]. While these preliminary tests were limited and not systematically evaluated, we observed frequent parsing errors such as missing commas or malformed coordinates. This experience suggested that a structured format like Geo-JSON could be easier to parse and work with, motivating its use in our experiments.

Birds exhibit higher predicted mean F1 scores than other groups (*p* = 0.024). While species occurring in Asia and Africa show a lower mean F1 in Figure D.13, these differences are not statistically significant after accounting for other factors in the regression model (*p* = 0.438) and (*p* = 0.513) respectively (Table D.8)). Similarly, no significant difference in performance is observed between `GPT-4o` and `Gemini 1.5 Pro` (*p* = 0.941). While no significant difference is observed, the limited number of species means small performance gaps cannot be ruled out.

### 3.3. Task 3: List Threatened Species in Region

Both models exhibit low mean precision, recall, and F1 with high standard deviations, indicating they miss a significant number of species and that the performance varies a lot between countries (Table 3). This difficulty stems from the models’ limitations when handling longer responses. In our case, the average response length is 2.3 species, with a maximum of 19 species identified across all countries. In countries like Indonesia, where up to 30 species need to be listed, this poses a particular challenge for both LLMs.

**Table 3:** Threatened Species. . The mean precision, recall, and F1 computed across countries for `Gemini 1.5 Pro` and `GPT-4o`. Geographic regions and countries without any known CR birds are not included in these scores.

| Model | Mean Precision | Mean Recall | Mean F1 |
| --- | --- | --- | --- |
| Gemini 1.5 Pro | 25.5 ± 35.7% | 18.0 ± 28.4% | 19.1 ± 26.9% |
| GPT-4o | **31.6**±39.3% | **20.1**±28.2% | **22.1**±27.7% |

The models demonstrate high recall and F1 scores in several countries with few critically endangered (CR) species (Figure 9). Accurately predicting just a few CR species in these nations leads to high scores. Many countries in southern Africa fall into this category. However, this performance is not uniformly high across all countries with low CR species counts and in many cases, such as Europe, F1 and recall are low. This inconsistency may arise because, when there are only a small number of correct species to predict, the model must be highly precise, making it challenging to generate outputs that align closely with the “ground truth”. The models are able to achieve high F1 and recall in Brazil, India, and Vietnam, even though they are countries with a higher number of CR bird species.

**Figure 9:**
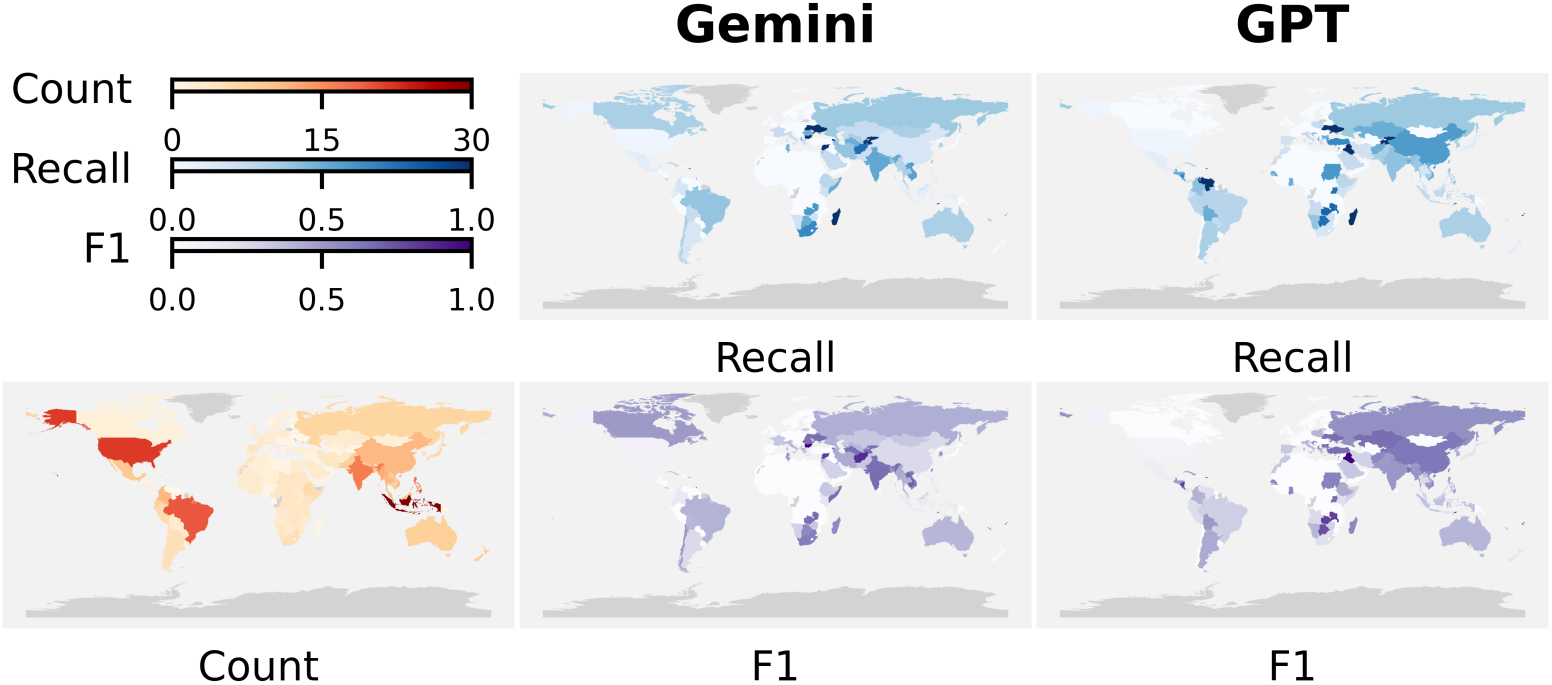
Task 3: Threatened Species. (Lower left) The number of critically endangered (CR) bird species across different countries, with darker red shades representing higher numbers. Grey indicates regions where data is unavailable. (Middle) The recall and F1 of `Gemini 1.5 Pro` when asked to list CR birds in different countries, with darker blue and purple shades representing higher recall and F1 respectively. (Right) The recall and F1 of `GPT-4o`.

### 3.4. Task 4: Threat Assessment

Neither model substantially outperforms a random guessing baseline when assessing threats (Table 4), where a scope and severity category is selected at random for each assessment. For scope classification, `GPT-4o` outperforms `Gemini 1.5 Pro` with an accuracy of 48.5% compared to `Gemini 1.5 Pro`’s 47.5%. In severity classification, `Gemini 1.5 Pro` shows a small advantage, with higher accuracy (28.3%) and cost-sensitive accuracy (71.7%) compared to `GPT-4o`, but both barely outperform the random baseline. Although the cost-sensitive accuracy indicates that the models often misclassify into adjacent categories, the results suggest that relying solely on zero-shot capabilities based on threat descriptions is insufficient.

**Table 4:**
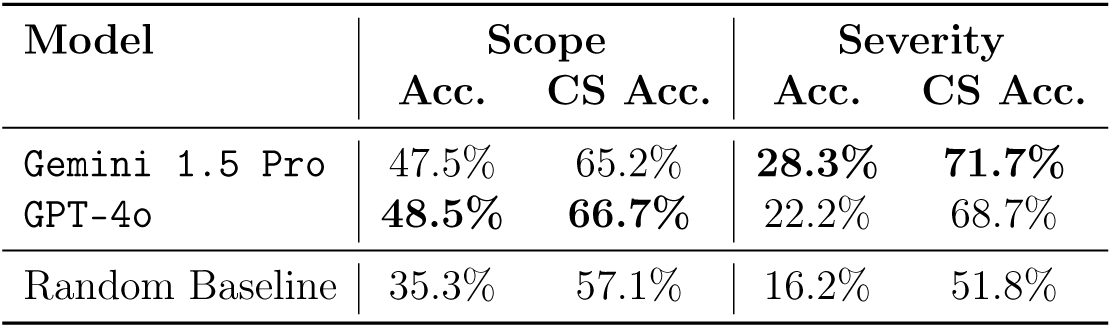
Threat Classification. . Accuracy (Acc.) and cost-sensitive accuracy (CS Acc.) for the scope and severity classification predictions made by `Gemini 1.5 Pro` and `GPT-4o` For the random guessing baseline, one scope and severity option is randomly selected per threat assessment; minor deviations from theoretical random accuracy (33% scope, 16.7% severity) arise due to stochastic variation, yielding the reported accuracies.

The models’ performance, which is only about ten percentage points higher than the random baseline, suggests that LLMs do not reliably perform this task. However, the cost-sensitive accuracy, which is higher for both models than the random baseline, indicates that when the models are incorrect, they tend to be closer to the correct answer than random guessing, though not substantially better. Both models tend to underestimate the scope by choosing “Minority” and tend to predict one of the medium severity options (Figure F.16).

### 3.5. Task 5: Trait Identification

`GPT-4o` consistently outperforms `Gemini 1.5 Pro` across both birds and mammals when predicting traits (Figure 10). For bird traits, `GPT-4o` achieves higher *R*^2^ values and lower Normalized Mean Absolute Error (NMAE) across all four traits, indicating more accurate predictions (Figure 10). Both models show lower performance for beak length, but `Gemini 1.5 Pro` additionally has low performance for the wing length predictions. Notably, an extreme overestimate for Turkey Vulture (*Cathartes aura*), with a prediction 1000 mm too large, results in a negative *R*^2^ score for wing length in `Gemini 1.5 Pro`.

**Figure 10:**
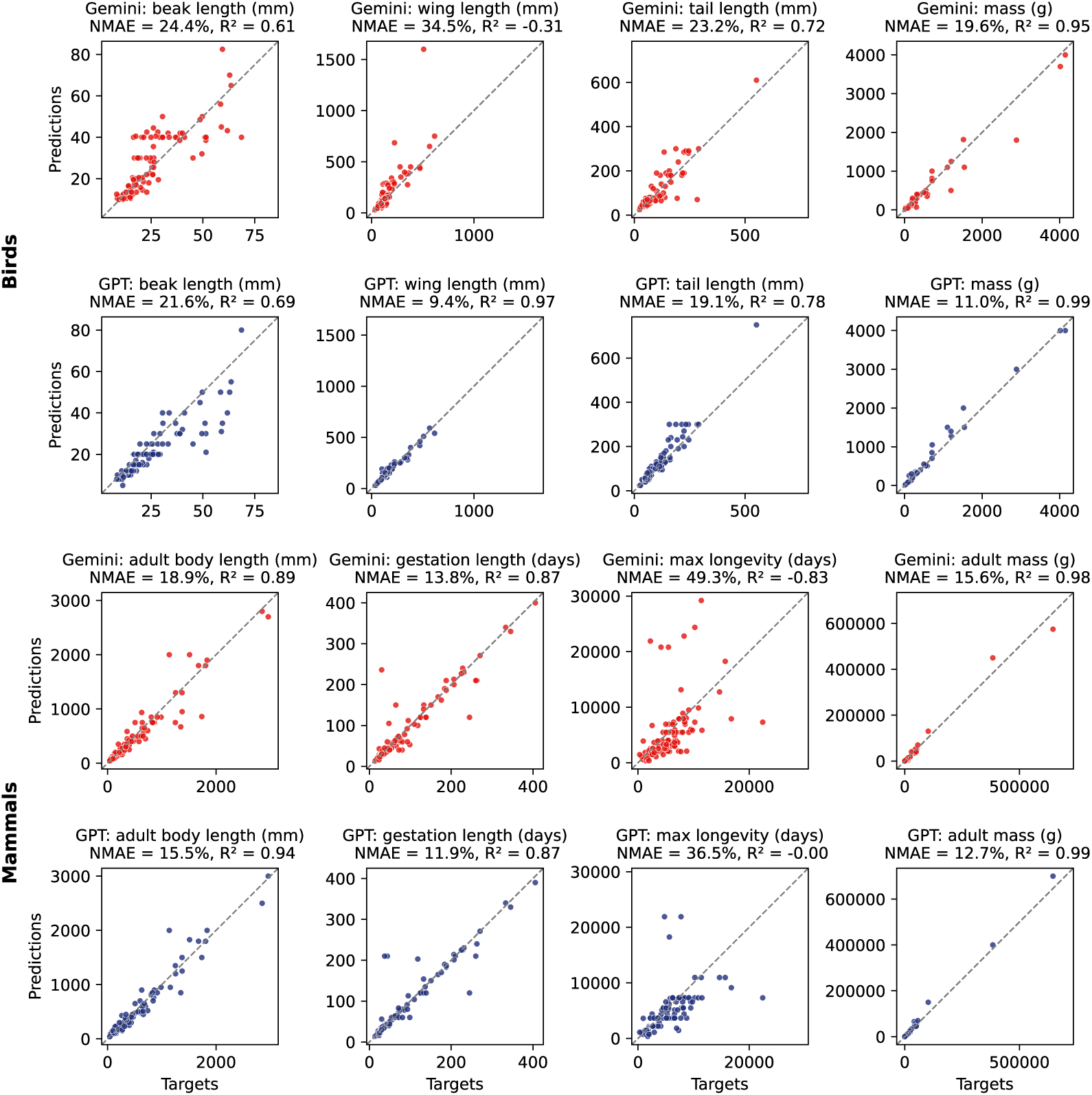
Trait Identification. Comparison of predictions between `Gemini 1.5 Pro` and `GPT-4o` for four bird traits: beak length, wing length, tail length, and mass, as well as four mammal traits: adult body length, gestation length, max longevity, and adult mass. Each panel shows a scatter plot of target values versus predictions with a reference line indicating a perfect prediction. NMAE (Normalized Mean Absolute Error) and *R*^2^ scores are reported for both models.

For mammals, `GPT-4o` similarly outperforms `Gemini 1.5 Pro` across all four traits. Both models have the lowest metrics for max longevity prediction, with `GPT-4o` achieving an *R*^2^ close to zero and `Gemini 1.5 Pro` yielding a negative *R*^2^.

In terms of output validity, `GPT-4o` also demonstrates greater reliability: we observe zero invalid responses out of 400 queries, compared to nine for `Gemini 1.5 Pro`. Failures in `Gemini 1.5 Pro` are due to inclusion of extra details, returning value ranges rather than single estimates, or null responses. Finally, while both models tend to make similar errors for mammal traits, greater divergence is observed for bird traits. In contrast, predictions for body mass, which is the only trait shared between birds and mammals in this experiment, show comparable error patterns across models and taxonomic groups, suggesting that the observed differences are more likely driven by the type and complexity of traits evaluated rather than taxonomy per se (Figure G.17).

## 4. Discussion

Large language models (LLMs) show promise for addressing ecological questions but their capabilities in this domain have previously been largely unexplored. In this study, we evaluated two leading commercial LLMs, `GPT-4o` and `Gemini 1.5 Pro`, across five ecological tasks, relying only on their internal parameters and prompt-provided information. Empirical evidence shows that LLMs retain substantial factual content in their parameters [49, 50]. Characterizing the scope and limits of this internal knowledge is therefore essential for understanding model capabilities and limitations in ecological applications. While our results are focused on two leading models, our benchmark provides a framework for evaluating other LLMs and for establishing baseline performance in the absence of retrieval.

Model performance shows substantial variability across task type, taxonomic group, and geography. The LLMs generally outperform simple baselines, with highest accuracy observed in presence–absence predictions, particularly at the country level (Task 1b). In contrast, performance declines for trait prediction tasks (Task 5) and range map generation (Task 2), which require precise numerical estimates and spatial coherence. The divergence between presence–absence and range map predictions highlights a gap in ecological knowledge: while the models can identify where a species occurs broadly, they struggle to produce maps that accurately reflect range limits. No statistically significant differences were observed between `GPT-4o` and `Gemini 1.5 Pro` for these geographic tasks, though our sample size of 100 species limits statistical power. Overall, performance appears to be shaped more by task structure than by model type, likely reflecting broader trends in training data coverage, where text-based and categorical information is more common than fine-grained geospatial or continuous trait data. These patterns align with recent ecological LLM studies, which show that accuracy in extracting species distribution data varies with source type and task complexity, and that performance in debunking wildlife misinformation depends on species group and context [28, 29]. Taken together with our results, these findings indicate that LLMs’ ecological knowledge is uneven, with particular weaknesses in tasks requiring more than factual recall, such as spatial inference or nuanced trait estimation.

The performance of LLMs in predicting species traits shows patterns that suggest a mix of memorization and inference. For example, when estimating traits like beak length, the models often produce values that cluster around a narrow range across species, forming horizontal bands in the plots. In contrast, predictions for traits such as wing length show greater variability and closer alignment with expert data. These differences may reflect cases where the model has been trained on sufficient data to memorize the pattern (e.g., for wing length), as compared to cases where the model needs to infer the value (e.g., beak length). Traits such as beak length and wing length are governed by known allometric relationships, which the model could leverage, though our trait-specific prompting likely limits the opportunity for such cross-trait inference. These patterns raise important questions about whether LLMs can draw ecological inferences (for example, apply allometric relationships) or primarily recalling associations encountered during training. An important avenue for improving trait prediction is therefore to move beyond purely text-based models. While this study focuses on LLMs, recent advances in multi-modal models that integrate text and images offer a promising direction. By grounding predictions in visual evidence, such models may reduce reliance on memorized averages and enable more specimen-specific inference, for example by estimating traits directly from photographs [51, 52].

Biases in training data present a core challenge for the application of LLMs in ecology. We observe that tasks involving classification of threats, such as scope and severity (Task 4), are especially difficult for the models, which often predict these incorrectly. This may be due to the relative rarity and complexity of such information in publicly available data. Even for human experts, such classifications are nuanced and context dependent. Errors in these predictions could misinform conservation decisions, for example by underestimating a species’ IUCN threat category, with downstream consequences for resource allocation and protection measures. Ethical considerations also include the amplification of biases, generation of plausible but false information, and potential misuse of sensitive ecological knowledge. Additionally, the computational cost of training and deploying LLMs is high [53], underscoring the need to balance benefits against environmental impacts.

While our evaluation was conducted in a “closed-book” setting, this design choice reflects an analytical goal rather than a recommended deployment mode. By largely withholding retrieval and external tools, we sought to isolate the ecological knowledge available to models to a combination of their parametric knowledge and the information provided in the prompt.

In practice, the most effective use of LLMs in ecology is likely to be within hybrid workflows, where models act as agents that query machine-readable biodiversity databases, invoke statistical and geospatial tools, and synthesize results across heterogeneous sources. In such settings, LLMs need not possess encyclopedic recall; however, they require a reliable substrate of background knowledge to correctly interpret retrieved information. Our results indicate that current models may be close to meeting this requirement: their robust performance on simpler tasks (significantly better than random overall) demonstrates a functional mixture of semantic understanding and knowledge of key ecological facts. This suggests that while in practice LLMs should likely rely on external tools for precision and timeliness, they possess the internal conceptual framework necessary to orchestrate those tools and support downstream analysis.

Conversely, we found that performance is lower for more complex tasks such as range mapping and threat classification, precisely the settings where evaluation is needed because errors can propagate into real conservation decisions. In these contexts, LLMs are better viewed as interfaces and synthesizers embedded within hybrid pipelines, rather than as autonomous sources of truth, with human-in-the-loop validation and explicit uncertainty estimates remaining essential. Importantly, our benchmark remains valuable with respect to this augmented setting: by establishing what models can and cannot do using only their parametric and prompt-conditioned knowledge, it provides a reference point against which retrieval-augmented systems can be evaluated, allowing future work to quantify the gains from leveraging external data and online retrieval. Moreover, because retrieval-augmented behaviors in current LLM systems are not always applied consistently across queries, closed-book evaluation provides a repeatable and controlled baseline of specific model versions.

Part of the discrepancy between LLM outputs and benchmark labels may reflect inherent uncertainty or aggregation in expert datasets, rather than model error alone. Future benchmarks should therefore aim to quantify such uncertainty while remaining ecologically meaningful and computationally reproducible. As LLM systems increasingly incorporate richer prompting strategies, in-context learning, and external tools, evaluation protocols must likewise evolve to capture how these factors shape performance. Prompt structure and context can substantially influence model behavior, and understanding their effects is essential for assessing reliability across ecological tasks. An important next step is the development of inference-based tasks, such as predicting outcomes of ecosystem interventions or imputing missing data from phylogenetic and spatial relationships, that go beyond factual recall. Systematically assessing performance across such tasks will enable more realistic measurement of ecological reasoning and guide improvements in model training, data integration, and workflow design.

## 5. Conclusions

We have presented an initial evaluation of LLMs in the ecological domain, which highlights both their current abilities, limitations, and points to their potential in the future. The commercial models evaluated show promise for tasks that involve recalling information, such as species presence predictions and trait estimation. However, their performance drops significantly in more complex tasks, such as generating accurate range maps or producing detailed threat classifications.

While these models could assist in accelerating certain conservation-related tasks and make ecological information more accessible, their limitations suggest that they are not yet ready for direct application in biodiversity decision making. Nonetheless, our new evaluation dataset represents a valuable resource that can be used to stimulate and facilitate future research, enabling advancements in the evaluation and application of LLMs to ecological and conservation challenges. Models that are more ecologically useful are likely to improve over time as techniques and training datasets evolve. However, it is important that we direct development towards ethically sound practices. Given the urgency of the biodiversity crisis, the benchmarking tasks introduced here can catalyze efficient and focused development of tools to help accelerate better conservation decision making.

## 6. Data availability

Code and data for reproducing our experiments is available at: https://github.com/filipgdorm/eco-llm.

## 7. Author contributions: CRediT

**FD**: Conceptualization, Methodology, Software, Validation, Formal Analysis, Investigation, Data Curation, Writing - Original Draft, Visualization. **JM**: Conceptualization, Methodology, Writing - Review & Editing. **DP**: Conceptualization, Methodology, Writing - Review & Editing. **MH**: Conceptualization, Methodology, Writing - Review & Editing. **OMA**: Conceptualization, Methodology, Resources, Writing - Review & Editing, Supervision, Project administration, Funding acquisition.

## 9. Declaration of Generative AI and AI-assisted Technologies in the Writing Process

During the preparation of this work the author(s) used Grammarly in order to improve the clarity and check spelling errors. After using this tool/service, the author(s) reviewed and edited the content as needed and take(s) full responsibility for the content of the published article.

## Appendix A. Datasets

The full list of species utilized for the tasks is available in the SPECIES.md file within the code repository.

## Appendix B. Evaluation Metrics

### F1 Score

The F1 score is computed in the overlap of cells between the LLM generated range map, and the expert range map.

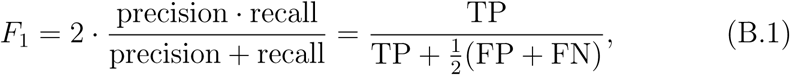

where TP denotes True Positives, FP False Positives, and FN False Negatives.

## Appendix C. Additional Details Task 1

**Figure C.11:**
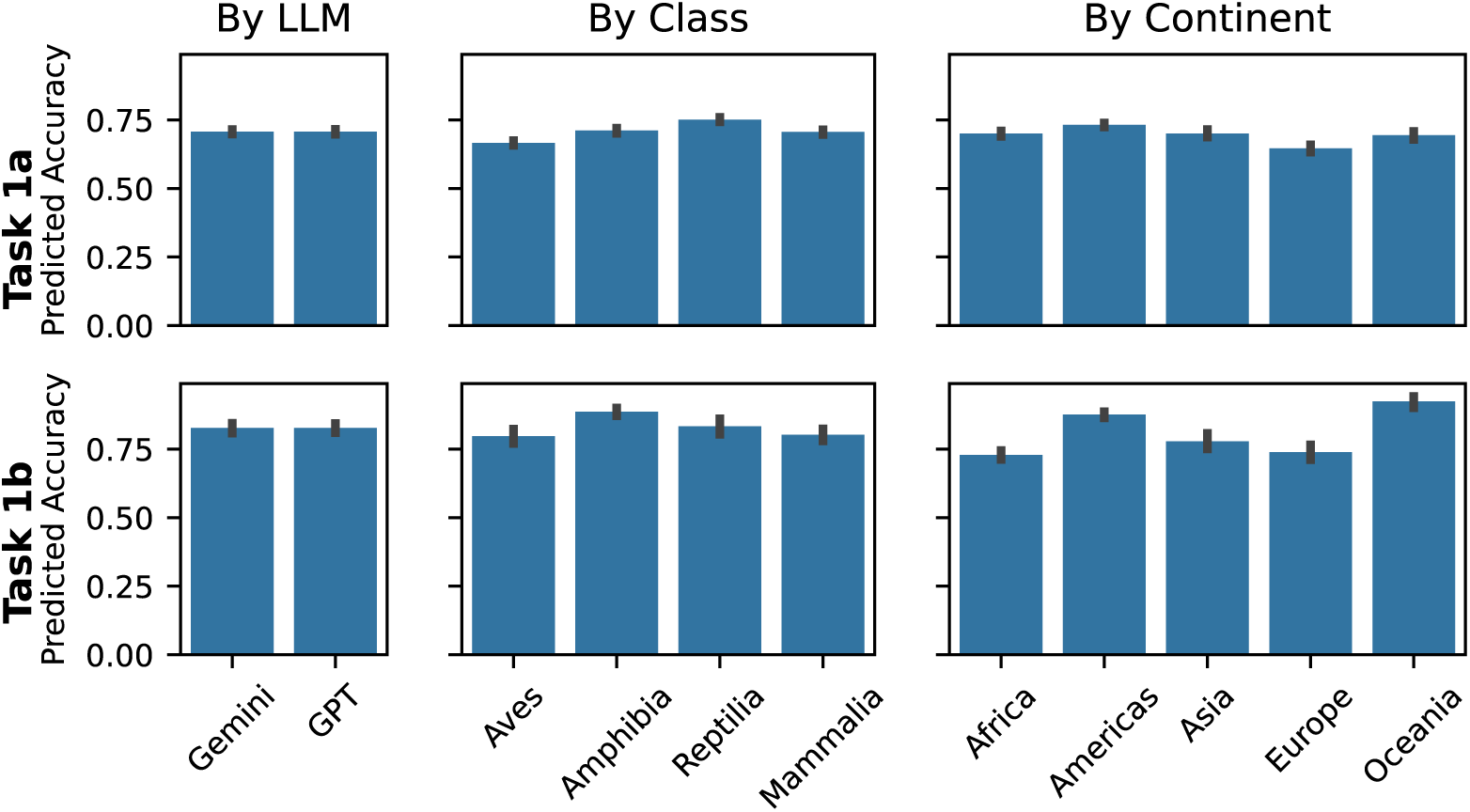
Predicted accuracy of species presence assessments for Task 1, grouped by language model (left), taxonomic class (middle), and species’ continental presence (right).

**Table C.5:**
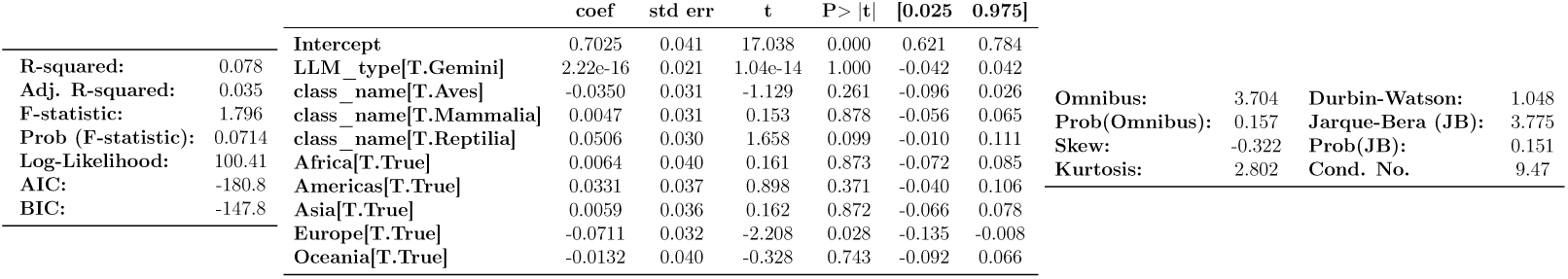
OLS Regression Results for Task 1a produced by the statsmodels package in Python.

**Table C.6:**
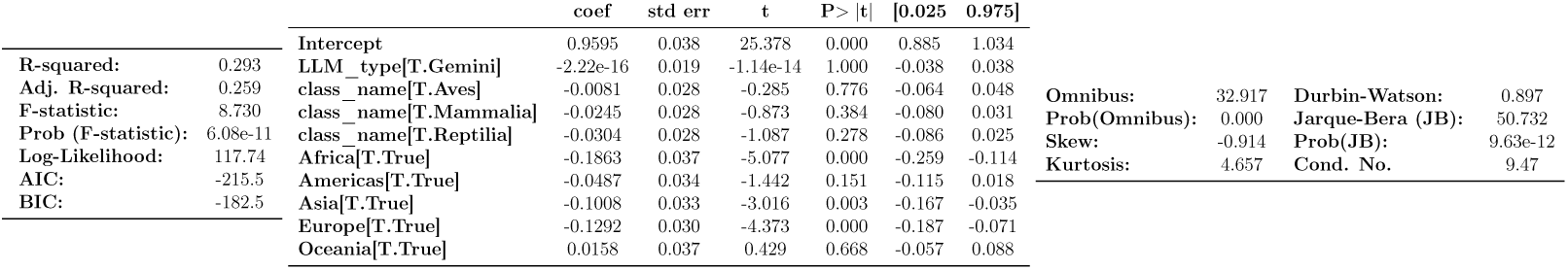
OLS Regression Results for Task 1b produced by the statsmodels package in Python.

### Appendix C.1. Difficulty of Absence Predictions

To assess whether LLMs perform differently on absences located near versus far from the true species range, we categorized all absence coordinates into *easy absences* (far from the known range) and *difficult absences* (near the range boundary). For each model, we computed the accuracy on presences, easy absences, difficult absences, as well as the overall and weighted accuracy.

**Table C.7:** Accuracy by presence and absence difficulty. Performance of each model across presences and absence categories, with overall accuracy combining all three groups.

| Model | Overall | Presence | Easy Absences | Difficult Absences |
| --- | --- | --- | --- | --- |
| Gemini 1.5 Pro | 0.708 | 0.644 | 0.981 | 0.459 |
| GPT-4o | 0.780 | 0.850 | 0.993 | 0.285 |

Both models achieve high accuracy on easy absences (those far from the species range), with over 98% correct classification. However, performance decreases for difficult absences near the range boundary, especially for `GPT-4o`, which correctly classifies only 28.5% of such points. Presence detection is moderate for `Gemini 1.5 Pro` (64.4%) and high for `GPT-4o` (85%), illustrating that fine-grained spatial reasoning near range limits remains challenging.

### Appendix C.2. Prediction Performance vs Range Size

We examine whether species range size affects Task 1a performance. Across species, we observe a small negative correlation between range size and model performance (Gemini: -0.16, GPT: -0.10), suggesting that model performance shows little dependence on range size (Figure C.12).

**Figure C.12:**
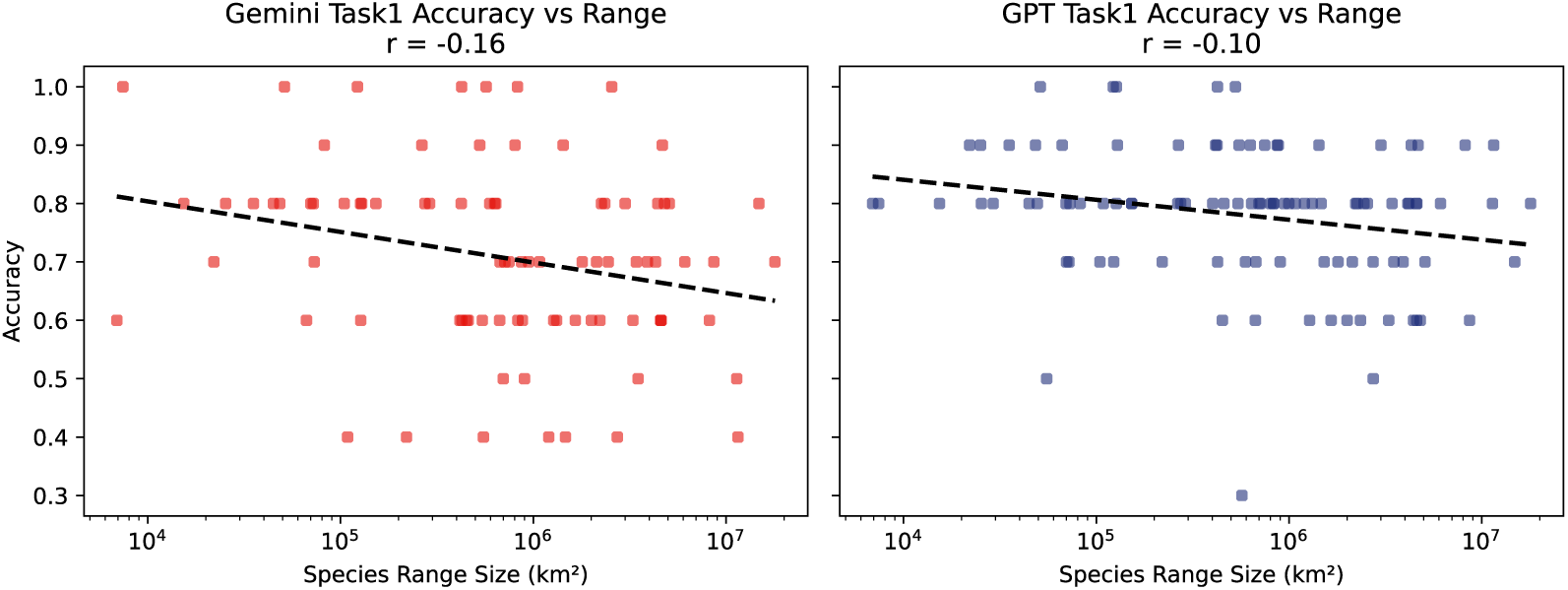
Task 1a performance versus species range size. Scatter plots show accuracy for each species, grouped by model: Gemini (left) and GPT (right). Trend lines indicate a small negative correlation between range size and performance (Gemini: *r* = -0.16, GPT: *r* = -0.10), suggesting that model performance is largely independent of range size under the fixed-presence/absence sampling scheme.

## Appendix D. Additional Details Task 2

**Table D.8:**
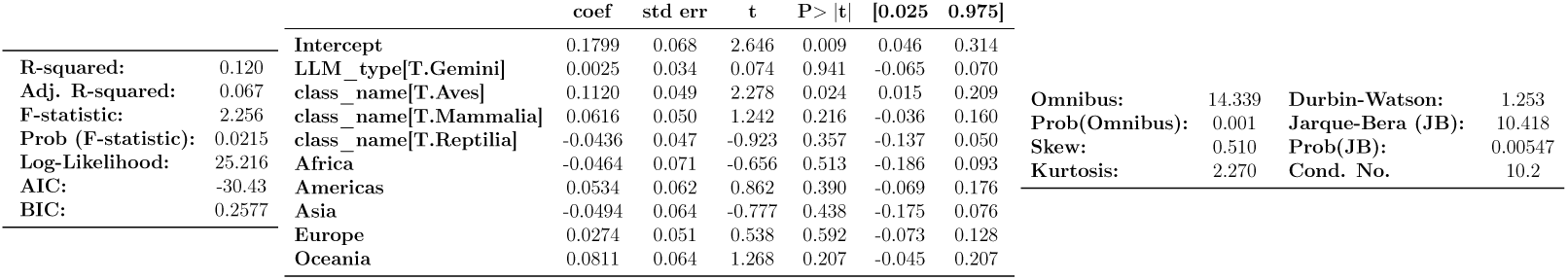
OLS Regression Results for Task 2 produced by the statsmodels package in Python.

## Appendix E. Consistency Between Geographical Tasks

We assess consistency between Task 1a (presence/absence classification) and Task 2 (species range estimation) using two complementary analyses.

**Figure D.13:**
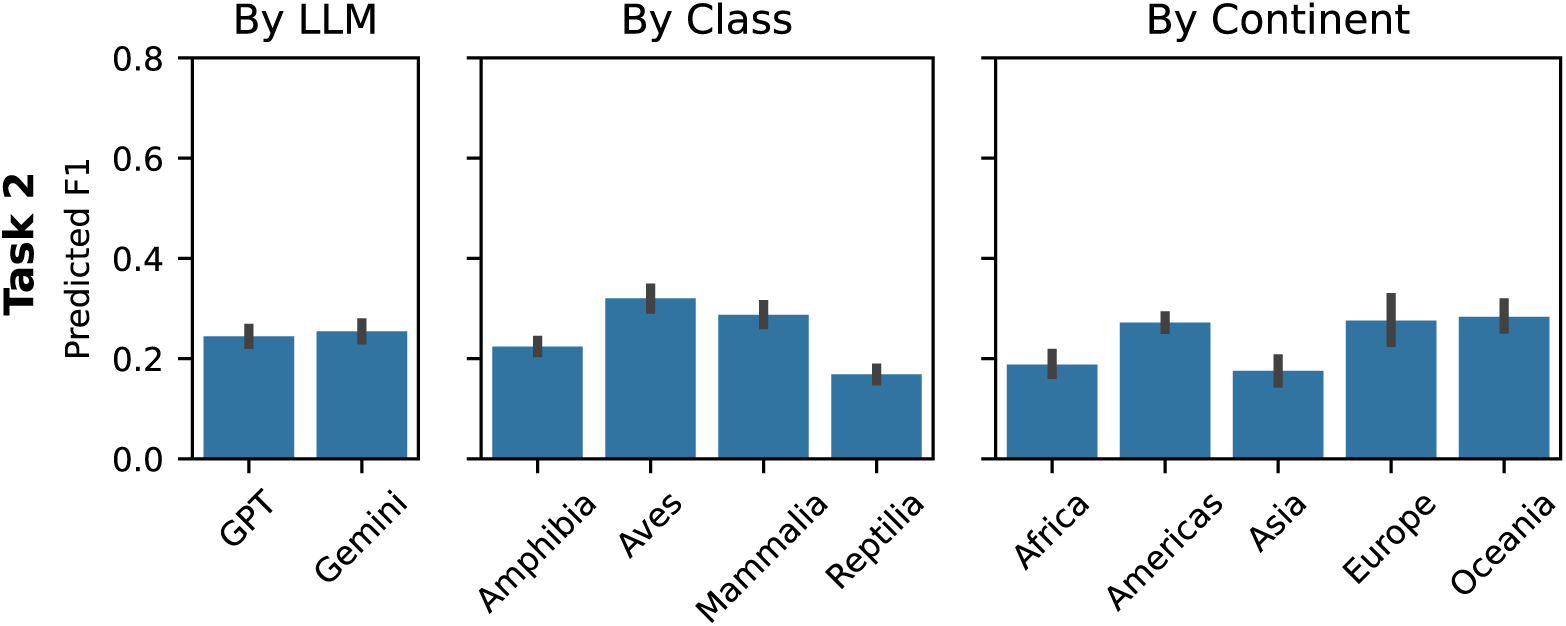
Predicted F1 of range maps for Task 2, grouped by language model (left), taxonomic class (middle), and species’ continental presence (right).

First, we examine the relationship between Task 1a and Task 2 outputs across species and models. Figure E.14 shows scatter plots comparing the species-level performance for both tasks. We observe no clear correlation, indicating that performance on Task 1a does not predict performance on Task 2.

**Figure E.14:**
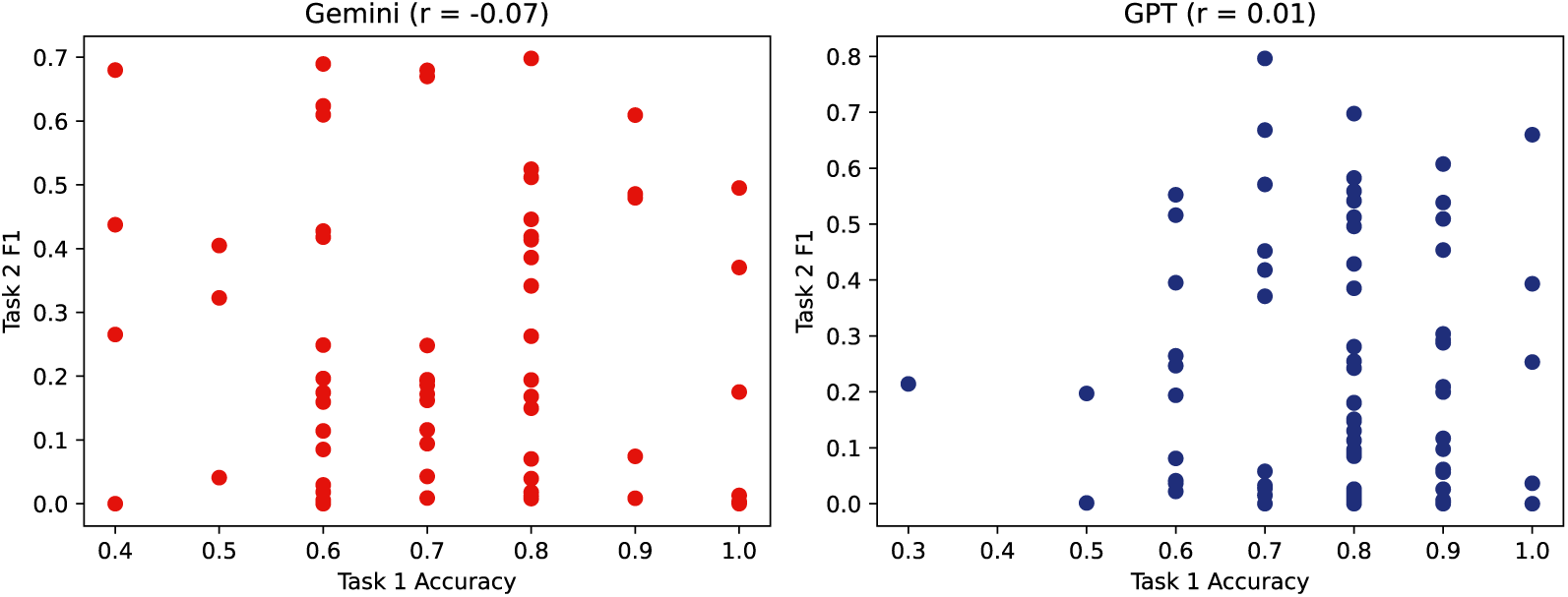
Consistency between Task 1 and Task 2. Scatter plots comparing Task 1 range estimates and Task 2 presence probabilities across species and models. No clear correlation is observed.

Second, we evaluate internal consistency by testing whether locations classified as presences in Task 1a fall within the range polygons generated by the same model in Task 2. For each species, ten queried locations are evaluated, and a consistency score is computed as the proportion of predicted presences lying within the inferred range.

We find low internal consistency for both models, with mean scores of 0.315 for `Gemini 1.5 Pro` and 0.274 for `GPT-4o`. This indicates that predicted presences frequently fall outside the model’s own inferred range, suggesting limited coherence between range-level and point-level predictions.

## Appendix F. Additional Details Task 4

**Figure F.15:**
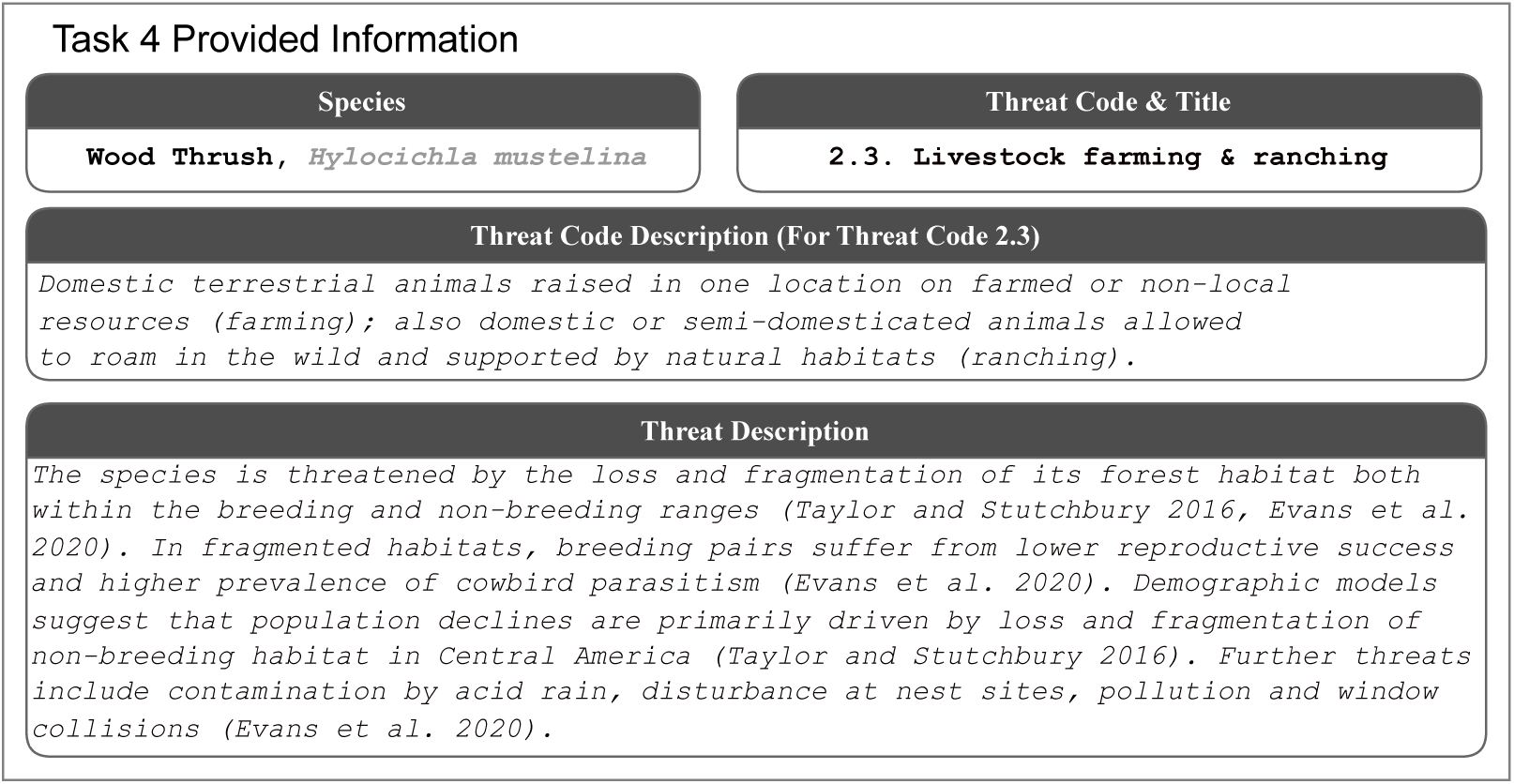
Example of the information provided to the model for Task 4 for the species Wood Thrush (*Hylocichla mustelina*). For each species, the model receives the species name, threat code, threat title, threat description. The information is sourced from the IUCN.

**Figure F.16:**
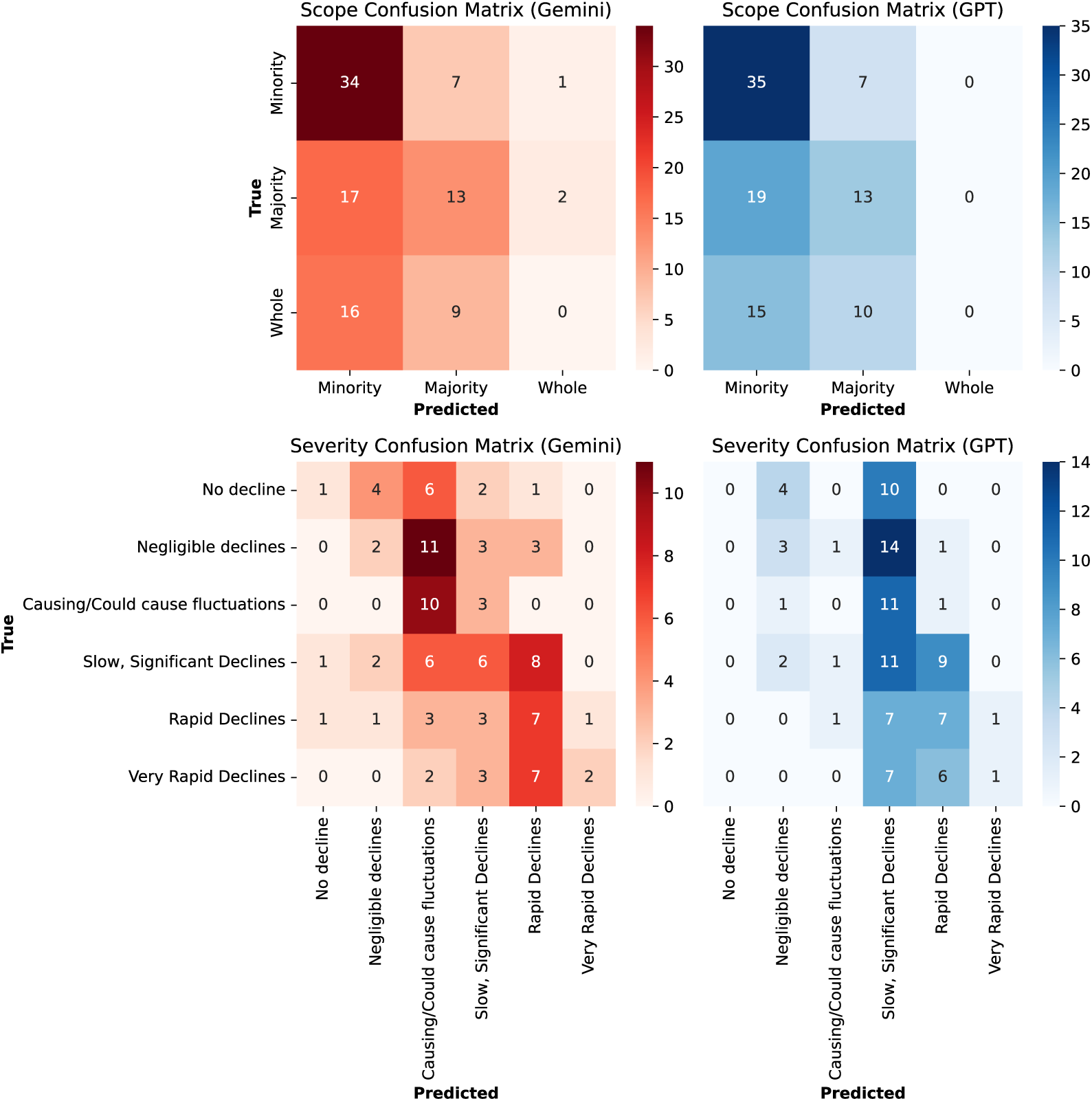
Task 4 Threat Assessment. The confusion matrices for Gemini and GPT for the scope and severity threat assessments.

## Appendix G. Additional Details Task 5

**Figure G.17:**
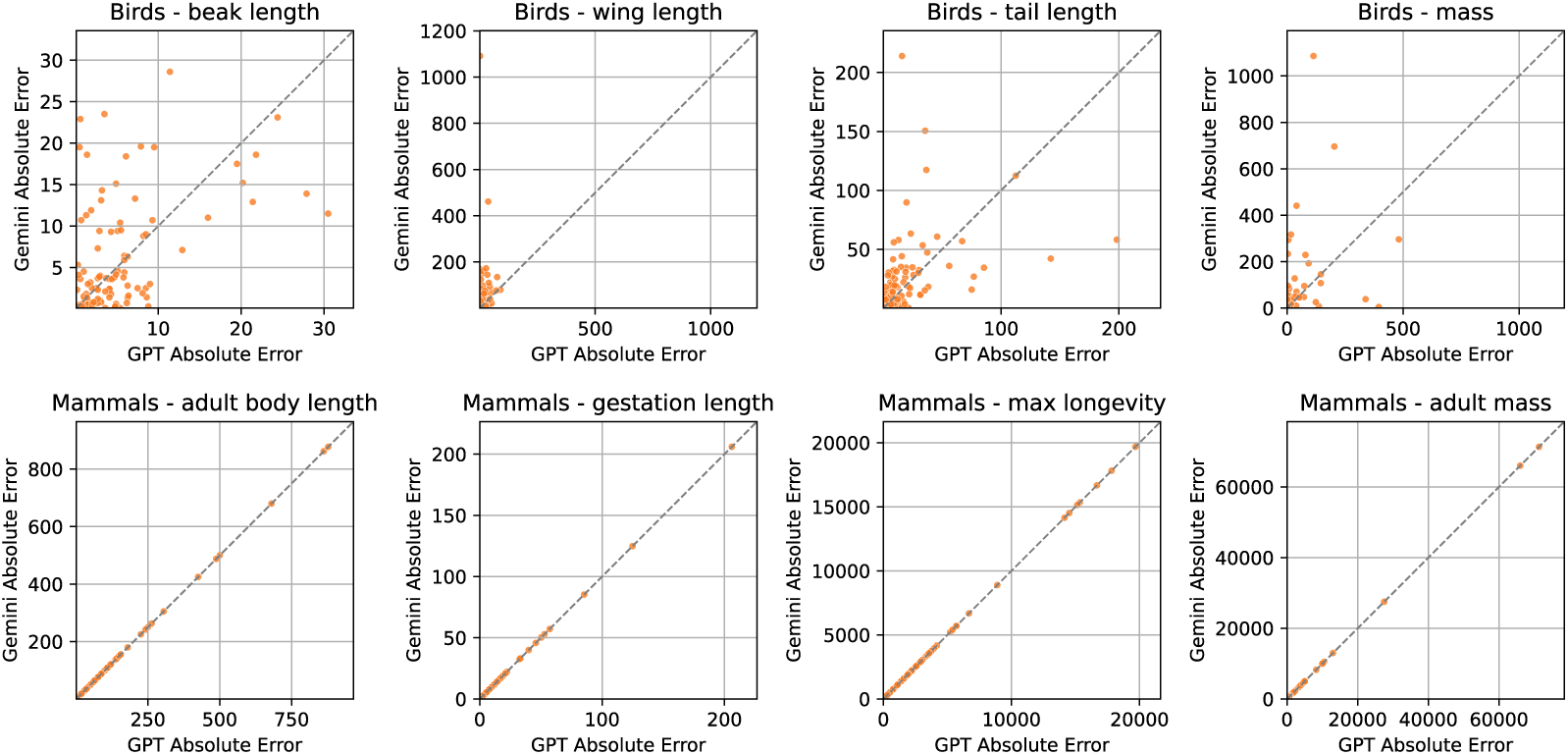
Task 5: Trait Identification. Scatter plot comparing species-level errors made by GPT (x-axis) and Gemini (y-axis) in Task 5. Each point represents a species, with its coordinates indicating the average error made by each model. Points along the diagonal line (y = x) reflect species for which both models perform similarly, while deviations indicate species where one model outperforms the other. This visualization helps identify whether the models struggle with the same species or exhibit complementary strengths.

